# Fiber Bragg grating based sensing system for non-destructive root phenotyping using ResNet prediction

**DOI:** 10.1101/2024.09.17.613457

**Authors:** Steven Binder, Kabir Hossain, Alexander Bucksch, Mable Fok

## Abstract

Real-time measurements of crop root architecture can overcome limitations faced by plant breeders when developing climate-resilient plants. Due to current measurement methods failing to continuously monitor root growth in a non-destructive and scalable fashion, we propose a first in-soil sensing system based on fiber Bragg gratings (FBG). The sensing system logs three-dimensional strain generated by a growing pseudo-root. Two ResNet models confirm the utility of in-soil FBG sensors by predicting pseudo-root width and depth with accuracies of 92% and 93%, respectively. To analyze model robustness, a preliminary experiment was performed where FBGs logged strain generated from a corn plant’s roots for 30 days. The models were then retrained on new data where they achieved accuracies of 98% and 96%, respectively. Our presented prototype has potential prospects to go beyond measuring root parameters and sense its surrounding soil environment.

## I. INTRODUCTION

BY 2050, our global population will reach upwards of 10 billion people and outpace the world’s production of agricultural resources [1]. Additionally, degraded soil conditions due to climate change and overuse of land will further limit crop production. Measurement of plant root architecture in a non-destructive, continuous fashion provides a promising means of improving nutrient uptake in crops [2]. Thus, providing a mitigation strategy for the adverse effects induced by climate change. With breeders developing more efficient crops, farmers will then be able to maximize the amount of food produced under limited water and nutrient availability.

The development of new root measurement tools is challenged by opaque soil; while the presence of highly conductive matter puts technologies based on electromagnetic waves at their resolution limit [3]. Existing methods for plant root phenotyping include destructive excavation using the Shovelomics protocol [4], X-ray tomography, magnetic resonance imaging (MRI) [5], and ground penetrating radar (GPR) [6]. While accurate, Shovelomics involves the destructive excavation of roots from soil before measurement [7], thus inferring biological meaning from only a part of the root system. GPR, X-ray, and MRI methods all measure non-destructively. However, they respectively have limitations such as low resolution measurements [7], observation of plants within size limited growth containers [8], and measurement distortions due to magnetic materials within the soil [9, 10]. In all cases, technicians are required to operate equipment during the measurement process, which is labor intensive and time consuming, preventing their deployment as autonomous sensing systems.

The advantage of fiber optic sensors, such as their immunity to electromagnetic interference, allows them to be deployed in a variety of environments. Additionally, their small size and passive construction insures they will not interfere with a growing plant [11]. Fiber optic sensors have previously been used for plant sensing. Such as a distributed fiber optic sensor that captures the strain of rice roots grown in washed Profile Greens Grade – an inorganic porous ceramic mixture [10]. While providing promising results, underground measurements using optical sensors underground still prove challenging due to the various soil parameters, such as temperature, moisture content, and compactness, that change over time. Previously, we have had promising results in measuring a deforming object within soil using fiber Bragg grating (FBG) sensors [12]. The FBG based setup provided optical power measurements related to the changing shape of an object in real time.

We propose a non-destructive root phenotyping sensing system utilizing fiber Bragg grating (FBG) sensors to log underground measurements to be used by residual neural network (ResNet) for root phenotype prediction. Our proposed system utilizes the reflected power of FBG wavelengths to minimize data size. To overcome the challenge of correlating raw sensor output with root parameters, our system utilizes two ResNet models for phenotyping root width and depth. Using ResNet ensures that gradient flow in the measurement data does not disappear during backpropagation learning steps [13]. After developing the models using data generated by a pseudo-root, they are then used in a preliminary experiment where they are retrained on real root growth data.

## II. Pseudo-Root Setup and ResNet Prediction

### A. Sensor and Pseudo-Root Deployment

In the presented experiment, our proposed sensing system detects and differentiates pseudo-root width classes of 1mm or 5mm and depth classes ranging from 0cm to 15cm with a resolution of 1.5cm. A reflected optical power-based measurement allows for a highly configurable system dependent on the number of plants to be monitored. Additionally, our ResNet model transforms the sensor’s complex output into root traits with known genetic underpinnings and relevance for plant breeding programs [14].

Fig. 1(a) illustrates the non-destructive sensing system, consisting of three FBG sensors with Bragg wavelengths of 1550.0nm, 1552.5nm, and 1553.3nm. Each FBG has its own wavelength-tunable laser diode and Thorlabs PM101 optical power meter for data collection. When strain is applied to the FBGs within soil, its Bragg wavelength elongates. Each laser is tuned to a wavelength that results in a power reading 3 dB lower than the sensors maximum possible reflected power. LabVIEW stores sensor output data as a text file for later use by the ResNet models. All experiments in this study were conducted at a constant room temperature and soil moisture content to eliminate any possible measurement distortions. Additionally, they were conducted using potting soil, which is the first demonstration of using FBGs for root sensing in real soil.

**Fig. 1.**
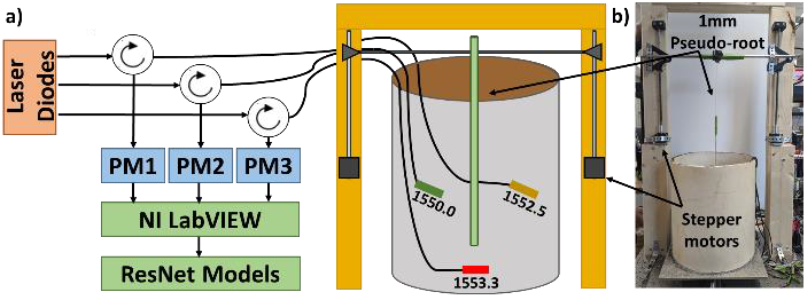
a) The non-destructive root phenotype sensing setup. b) The physical robotic root setup.

The two side FBGs, with wavelengths of 1550.0nm and 1552.5nm, were 8.5cm below the soil surface and 2cm offset from the insertion point. The bottom FBG, with a Bragg wavelength of 1553.3nm, was directly centered under the insertion point, 17cm below the soil surface. All FBGs were provided with a 2cm buffer so that they would not encounter the pseudo-root. Each FBG was oriented such that they were perpendicular for maximum strain coupling.

To efficiently generate large quantities of pseudo-growth data, we built a robotic system as shown in Fig. 1(b). The robot allows for the timed insertion of a pseudo-root into soil until a specific depth is reached. Controlling insertion time allows for a better correlation between sensor power change and pseudo-root parameters when training the ResNet models. Leading to higher model accuracy when tested on new data.

Two pseudo-roots with diameters of 1mm and 5mm are inserted into soil ten times to a depth of 15cm over 11 minutes to generate two datasets. Due to the complex nature of the FBGs output, seen in Fig. 2, two separate data preprocessing methods are introduced.

**Fig. 2.**
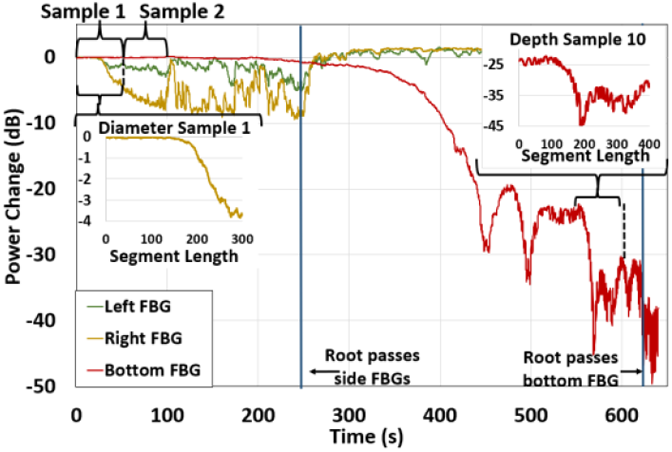
The induced FBG power change by the 5 mm pseudo-root and the sample processing used for each trial.

### B. ResNet Development and Results

Samples within the width and depth datasets are set to have fixed segment lengths of 300 and 400 data points and yielded a total of 3,228 and 300 samples, respectively. The width samples were generated using data from the two side FBGs and depth sample creation used only the bottom FBG output. Each width sample contains a single root diameter, while a depth sample represents a single depth range between 0cm to 15cm in 1.5cm increments. Fig. 2 provides an example of processed samples with fixed segment lengths. Furthermore, samples of both datasets were augmented with a sliding window approach. The sliding window overlapped each original segment ranging in coverage from 0% to 83.33% to help enrich the final datasets. The proposed ResNet model starts with a 2×2 convolution layer (16 filters), followed by batch normalization, ReLU activation, and 2×2 max pooling. This is followed by 25 residual blocks, each with two 5×5 convolution layers (16 filters), batch normalization, and ReLU activation.

The output layer includes neurons corresponding to the number of classes: two for width and ten for depth (pseudo-root), and four for width and five for depth (real corn roots). Our model implementations are available as Python code using the TensorFlow module [15] and can be found in reference [16].

Processed datasets were partitioned using an 80/20 split for training and testing. During the training phase, we applied dropout regularization to our ResNet model, randomly dropping units and their connections to prevent overfitting [17]. Additionally, we utilized data augmentation techniques, such as sliding window methods, to increase the size of our datasets. These protocols helped improve the model’s generalization capability and mitigate the risk of overfitting.

During testing, we evaluated model accuracy, precision, True Positive Rate (TPR), and False Positive Rate (FPR) as seen in Table 1. Seen in Fig. 3(a) is an example of the width model’s predictive results achieved from test data. With a TPR of 93%, the model captured root diameters reliably. However, the slightly higher FPR of 9.2% suggests misclassification of negative instances as positives. This is likely caused by the model’s sensitivity towards achieving a higher recall as seen by its 93% accuracy and 90% precision scores. By maintaining a high recall, this ensures that most true positives are identified. Similarities in the FBG signals provided by the different diameter classes may arise due to shared environmental factors, such as soil compactness or moisture levels. Nonetheless, the model’s precision and recall demonstrate its effectiveness in accurate predictions.

**TABLE I.** Performance metrics of both models on pseudo-root data.

| Model | Accuracy | Precision | TPR | FPR |
| --- | --- | --- | --- | --- |
| Width | 92% | 90% | 93% | 9.2% |
| Depth | 93% | 94% | 93% | 0.74% |

**Fig. 3.**
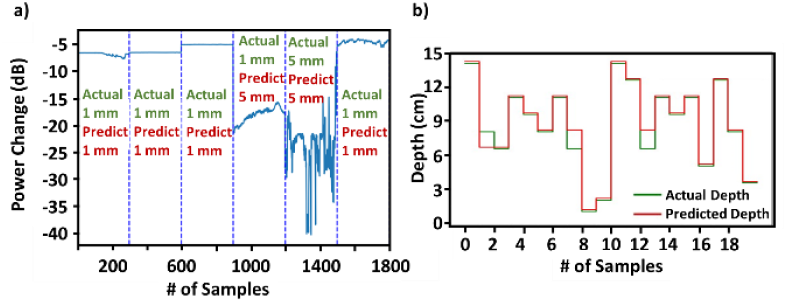
The predictive output of the (a) diameter and (b) depth models when tested on novel data.

The depth model’s precision signifies correct depth predictions 94% of the time. Additionally, its high TPR of 93% indicates the model’s reliability in correctly predicting the current depth level. The low FPR of 0.74% proves the model rarely produces incorrect predictions. Fig. 3(b) shows an example of the results from our depth prediction model, where the red trace shows the predicted class provided by our ResNet model and the green trace shows the sample’s actual class.

Fig. 4(a) visualizes our diameter model’s Area Under Curve (AUC) which highlights its performance and the tradeoff that occurs between TPR and FPR. The AUC value is a quantitative measure of overall performance; a higher AUC indicates better discrimination between positive and negative instances. Seen in Fig. 4(b) is our depth model’s confusion matrix produced after testing. The diagonal elements represent correct predictions for each depth level, reflecting accurate depth level identifications. Off-diagonal elements indicate instances of misclassification. For example, Depth Level (DL) 3 contained 5 samples for model testing. Out of these, 4 were correctly identified while 1 was misclassified as indicated by the off-diagonal element.

**Fig. 4.**
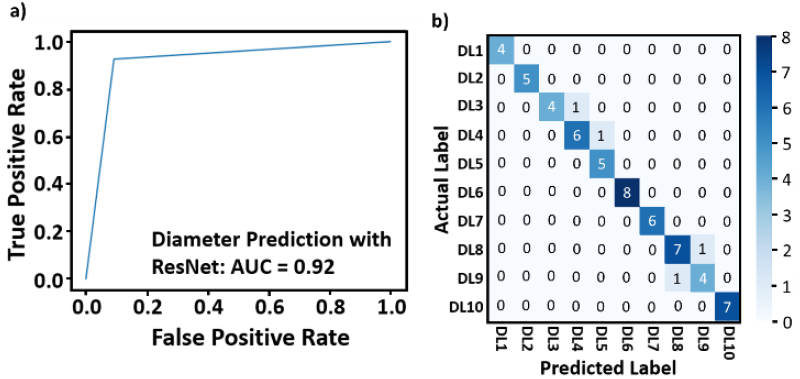
(a) The diameter model’s AUC curve. (b) The depth model’s confusion matrix.

## III. ResNet Performance on Real Root Data

### A. Sensor Deployment

To validate model performance on real roots, three FBGs were embedded within soil to measure the growth of a corn seedling for a preliminary demonstration. A 1550.0nm and 1552.5nm FBG was placed at the same depth as the seed but 2cm to the left and right. A 1553.3nm FBG was placed directly below the seed 2cm as seen in Fig. 5. An additional 1547.0nm FBG was covered with a protective casing and placed away from the seed to serve as a reference sensor. Its output was later used in removing soil temperature and moisture induced variations. Unlike in the previous setup, an optical spectrum analyzer (OSA) was used to measure each FBGs wavelength shift. Doing so provided greater data insight and flexibility for viewing the root’s growth overtime. Python was used to extract the reflected power of specific wavelengths to be used in model retraining. FBG measurements were taken for a total of 30 days after seed germination.

**Fig. 5.**
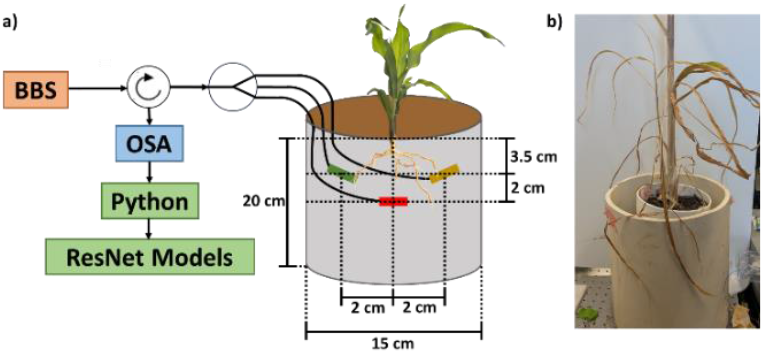
(a) The experimental setup used for measuring corn root growth. (b) The corn 30 days after germination.

### B. ResNet Retraining and Results

Due to the small dataset size obtained from this preliminary experiment, robust augmentation techniques were employed for data enrichment and enhancing our model’s generalization capabilities. Before being augmented with noise addition [18], scaling [19], and time shifting [20], the dataset was split into training and testing sets. Unlike in the pseudo-root experiment, a 70:30 split was used to ensure the test size was not smaller than the number of predicted classes. Studies suggest that this split for training and testing will still yield optimal results [21, 22].

While data was collected for a total of 30 days, only 12 were used in width and 15 for depth prediction as the roots grew past the sensors after this time. As a result, the width dataset contained 960 samples for training and 40 for testing. Each sample was provided a class of 0-3.9, 4-5.9, 6-7.9, or 8-10.0cm noting the width of the corn’s root system. The depth dataset contained 1200 samples for training and 50 for testing with class labels of 0-1.9, 2-7.9, 8-9.9, 10-11.9, and 12-14.0cm. The class labels were determined through studies on average corn root growth in unstressed conditions [23].

Table 2 provides both models’ performance metrics on the test data. When trained on actual root data, both ResNet models provide accurate predictions regarding root width and depth. As seen in the width (Fig. 6(a)) and depth (Fig. 6(b)) confusion matrices, the model’s only struggle with predicting roots as they grow larger. However, this inaccuracy may be attributed to sensing the strain generated by the growth of several smaller offshoots near the sensors or the roots growing past the FBGs.

**TABLE II.** Performance metrics of both models on real root data.

| Model | Accuracy | Precision | TPR | FPR |
| --- | --- | --- | --- | --- |
| Width | 98% | 98% | 97% | 1.0% |
| Depth | 96% | 96% | 96% | 1.0% |

**Fig. 6.**
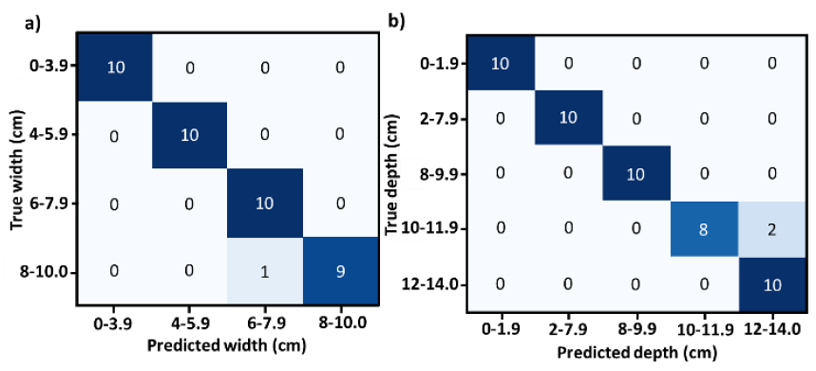
(a) The width and (b) the depth model’s confusion matrix.

## IV. Conclusion

In summary, we propose a non-destructive and continuous approach for quantifying root growth in real-time. Three FBGs embedded in soil paired with ResNet deep learning models, reliably predicts a pseudo-root’s width and depth with high accuracy. When the ResNet models are retrained on growth measurements of a real corn root, the models successfully predict its growth overtime. A benefit of the presented sensing configuration is that additional FBGs can be added to increase sensitivity and resolution.

Performance metrics of the two ResNet models suggest only a small chance of misclassification when presented with new data. Both ResNet models convert sensor data into root trait measurements with known genetic underpinnings. Thus, our measurements will be useful to breeders for developing climate-resilient plants as we progress toward training the models on actual root growth data. While performing accurately, we do recognize the relatively small dataset size in this preliminary study. With base models created, our next steps include the growth of several corn seedlings under controlled environmental conditions with more FBGs to sense strain generated by different growth phases. Using new datasets generated from crop plants, the model’s adaptability and generalization will be enhanced.

## References

[1] A. Bucksch et al., “Morphological Plant Modeling: Unleashing Geometric and Topological Potential within the Plant Sciences,” (in English), Front Plant Sci, vol. 8, p. 900, Jun 9 2017, doi: 10.3389/fpls.2017.00900.

[2] J. P. Lynch, “Roots of the second green revolution,” (in English), Australian Journal of Botany, vol. 55, no. 5, pp. 493–512, 2007, doi: 10.1071/Bt06118.

[3] R. W. T. Pena, P. A. C. Oliva, and F. A. Abrunhosa, “Application of the Ground Penetrating Radar (GPR) and Electromagnetic (EM34-3) Geophysical Tools and Sedimentology for the Evaluation of the Subsurface of Sites Earmarked for Aquaculture Ponds in the Amazon Region of Northern Brazil,” (in English), Applied Sciences-Basel, vol. 13, no. 19, p. 11107, Oct 2023, doi: 10.3390/app131911107.

[4] J. Kengkanna, P. Jakaew, S. Amawan, N. Busener, A. Bucksch, and P. Saengwilai, “Phenotypic variation of cassava root traits and their responses to drought,” (in English), Applications in Plant Sciences, vol. 7, no. 4, p. e01238, Apr 2019, doi: 10.1002/aps3.1238.

[5] R. Metzner et al., “Direct comparison of MRI and X-ray CT technologies for 3D imaging of root systems in soil: potential and challenges for root trait quantification,” Plant Methods, vol. 11, no. 1, p. 17, 2015, doi: 10.1186/s13007-015-0060-z.

[6] X. W. Liu, X. J. Dong, and D. I. Leskovar, “Ground penetrating radar for underground sensing in agriculture: a review,” (in English), International Agrophysics, vol. 30, no. 4, pp. 533–543, Oct 2016, doi: 10.1515/intag-2016-0010.

[7] A. Li, L. Zhu, W. Xu, L. Liu, and G. Teng, “Recent advances in methods for in situ root phenotyping,” PeerJ, vol. 2021, p. e13638, 03/11/2021 2022, doi: 10.34133/2021/8696571.

[8] H. Poorter, B. H. J, D. van Dusschoten, J. Climent, and J. A. Postma, “Pot size matters: a meta-analysis of the effects of rooting volume on plant growth,” Funct Plant Biol, vol. 39, no. 11, pp. 839–850, Nov 2012, doi: 10.1071/FP12049.

[9] G. C. Bagnall et al., “Design and demonstration of a low-field magnetic resonance imaging rhizotron for in-field imaging of energy sorghum roots,” The Plant Phenome Journal, vol. 5, no. 1, p. e20038, 2022, doi: 10.1002/ppj2.20038.

[10] M. Tei, E. Barbieri, F. Soma, Y. Uga, and Y. Kawahito, “Agritech imaging of underground plant root growth using a distributed fiber optic sensor,” in Optical Fibers and Sensors for Medical Diagnostics, Treatment and Environmental Applications XXII, San Francisco, CA, 2022 2022, vol. 11953: SPIE, pp. 98–102, doi: 10.1117/12.2606308.

[11] C. Pendao and I. Silva, “Optical Fiber Sensors and Sensing Networks: Overview of the Main Principles and Applications,” Sensors (Basel), vol. 22, no. 19, p. 7554, Oct 5 2022, doi: 10.3390/s22197554.

[12] S. Binder, M. Yang, V. Qiu, A. Bucksch, and M. Fok, “Non-destructive measurements of root traits and their soil-water environment using Fiber Bragg Grating-based fiber optic sensors,” Authorea Preprints, 2022, doi: 10.1002/essoar.10508796.1.

[13] Y. Bengio, P. Simard, and P. Frasconi, “Learning Long-Term Dependencies with Gradient Descent Is Difficult,” (in English), Ieee Transactions on Neural Networks, vol. 5, no. 2, pp. 157–166, Mar 1994, doi: Doi 10.1109/72.279181.

[14] W. Ren et al., “Genome-wide dissection of changes in maize root system architecture during modern breeding,” Nat Plants, vol. 8, no. 12, pp. 1408–1422, Dec 2022, doi: 10.1038/s41477-022-01274-z.

[15] V. Ivan, Python deep learning: exploring deep learning techniques and neural network architectures with PyTorch, Keras, and TensorFlow. Packt Publishing, 2019.

[16] K. Hossain. “Root-Diameter-Prediction-with-Residual-Neural-Network.” https://wave.engr.uga.edu/code/

[17] N. Srivastava, G. Hinton, A. Krizhevsky, I. Sutskever, and R. Salakhutdinov, “Dropout: a simple way to prevent neural networks from overfitting,” The journal of machine learning research, vol. 15, no. 1, pp. 1929–1958, 2014.

[18] M. Sáiz-Abajo, B.-H. Mevik, V. Segtnan, and T. Næs, “Ensemble methods and data augmentation by noise addition applied to the analysis of spectroscopic data,” Analytica chimica acta, vol. 533, no. 2, pp. 147–159, 2005.

[19] K. He, X. Zhang, S. Ren, and J. Sun, “Deep residual learning for image recognition,” in Proceedings of the IEEE conference on computer vision and pattern recognition, 2016, pp. 770–778.

[20] M. B. Er and I. B. Aydilek, “Music emotion recognition by using chroma spectrogram and deep visual features,” International Journal of Computational Intelligence Systems, vol. 12, no. 2, pp. 1622–1634, 2019.

[21] A. Gholamy, V. Kreinovich, and O. Kosheleva, “Why 70/30 or 80/20 relation between training and testing sets: A pedagogical explanation,” Int. J. Intell. Technol. Appl. Stat, vol. 11, no. 2, pp. 105–111, 2018.

[22] V. R. Joseph, “Optimal ratio for data splitting,” Statistical Analysis and Data Mining: The ASA Data Science Journal, vol. 15, no. 4, pp. 531–538, 2022.

[23] F. Hochholdinger, K. Woll, M. Sauer, and D. Dembinsky, “Genetic dissection of root formation in maize (Zea mays) reveals root-type specific developmental programmes,” Annals of Botany, vol. 93, no. 4, pp. 359–368, 2004.

